# Identification of sample mix-ups and mixtures in microbiome data in Diversity Outbred mice

**DOI:** 10.1101/529040

**Authors:** Alexandra K. Lobo, Lindsay L. Traeger, Mark P. Keller, Alan D. Attie, Federico E. Rey, Karl W. Broman

**Author notes:** **Corresponding author:** Karl W Broman, Department of Biostatistics and Medical Informatics, University of Wisconsin–Madison, 2126 Genetics-Biotechnology Center, 425 Henry Mall, Madison, WI 53706.

## Abstract

In a Diversity Outbred mouse project with genotype data on 500 mice, including 297 with microbiome data, we identified three sets of sample mix-ups (two pairs and one trio) as well as at least 15 microbiome samples that appear to be mixtures of pairs of mice. The microbiome data consisted of shotgun sequencing reads from fecal DNA, used to characterize the gut microbial communities present in these mice. These sequence reads included sufficient reads derived from the host mouse to identify the individual. A number of microbiome samples appeared to contain a mixture of DNA from two mice. We describe a method for identifying sample mix-ups in such microbiome data, as well as a method for evaluating sample mixtures in this context.

## Introduction

Sample mix-ups and other sample mislabelings interfere with our ability to map the genetic loci affecting complex phenotypes, as the mis-alignment of genotype and phenotype will weaken associations, reduce power, and give biased estimates of locus effects.

In traditional genetic studies, one has limited ability to detect sample mix-ups. Inconsistencies between subjects’ sex and X chromosome genotypes may reveal some problems, but one is blind to most errors. However, in studies that include genome-wide gene expression data, there is an opportunity to not just identify but also correct sample mix-ups (Westra *et al.* 2011; Lynch *et al.* 2012; Broman *et al.* 2015), as strong local expression quantitative trait loci (eQTL) can be used to identify individuals.

Studies to identify host loci that affect microbiome population composition could similarly be affected by sample mix-ups. If the microbiome analysis focuses on a single locus (such as 16S ribosomal RNA), one would have little ability to detect mix-ups. But when applying shotgun sequencing to microbiome samples to obtain metagenomic data, there may be sufficient sequence reads from the host to identify the individual. Fecal samples contain a variable fraction of host DNA that comes from intestinal epithelial cells. These cells are renewed every 4–5 days (Eastwood 1977), and dead cell slough off into the lumen and are eliminated in feces.

In a Diversity Outbred (DO) mouse project with genotype data on 500 animals (Keller *et al.* 2018), we have metagenomic sequencing data collected from fecal DNA of 297 mice. DO mice (Churchill *et al.* 2012; Svenson *et al.* 2012) are an advanced intercross population derived from eight founder strains (the same founders as the Collaborative Cross; Churchill *et al.* 2004). The metagenomic sequence data include sufficient reads derived from the host to identify the individual. We describe a method for identifying sample mix-ups in such microbiome data, as well as a method for evaluating sample mixtures in this context. We identified three sets of sample mix-ups (two pairs and one trio) as well as at least 15 microbiome samples that appeared to be mixtures of pairs of mice.

## Material and Methods

This study was part of a larger study (see Keller *et al.* 2018) that involved 500 Diversity Outbred (DO) mice that were obtained in five batches of 100 mice each, from generations 17, 18, 19, 21, and 23. The mice were obtained from the Jackson Laboratories (stock no. 009376) at 4 weeks of age, were maintained on a high fat/high sucrose Western-style diet, and were singly housed.

### Microbiome data

DNA was extracted from the feces of three of the five batches of DO mice (1, 2, and 4). DNA was isolated as described in Turnbaugh *et al.* (2009) and Kreznar *et al.* (2017). Libraries were prepared for Illumina sequencing according to the methods laid out in McNulty *et al.* (2011) with the following alteration: in the gel purification stage, DNA fragments were isolated at ∼450 bp instead of 200 bp. Paired-end sequencing (2×125) was performed using both MiSeq and Illumina HiSeq 2500 technologies, to allow for testing libraries before Hiseq runs, and to obtain additional sequences from under-sequenced samples. Raw reads were demultiplexed using the Fastx Toolkit (version 0.0.13; Hannon 2010), `fastx_barcode_splitter.pl` with `partial 2` and `mismatch 2`. Demultiplexed reads from the same samples that occurred on multiple lanes or sequencing platforms were concatenated into one forward and one reverse read file. Reads were then trimmed to remove barcodes (`fastx_trimmer −f 9 −Q 33`) and for read quality (`fastq_quality_trimmer −t 20 −l 30 −Q33`). Trimmed reads files were re-paired (with unpaired reads removed) using custom perl scripts. Reads were mapped to the mouse genome (build GRCm38/mm10) using bowtie2 (version 2.2.7; Langmead and Salzberg 2012) with default settings. Host-derived reads were identified using samtools (version 1.3; Li *et al.* 2009). Three metagenomic samples did not yield DNA and were excluded from further analysis (DO-148, DO-182, and DO-202).

### SNP genotypes

Mouse genotyping was performed on tail biopsies as described in Svenson *et al.* (2012), using the third-generation Mouse Universal Genotyping Array (GigaMUGA; Morgan *et al.* 2016) at Neogen (Lincoln, NE). The GigaMUGA array includes 143,259 SNPs. We focused on 110,143 autosomal SNPs that were placed in quality tiers 1 or 2 (out of 4) in Morgan *et al.* (2016).

We used a hidden Markov model (HMM) to calculate the probability of each of the 36 possible diplotypes (that is, pairs of founder haplotypes) along each chromosome in each DO mouse, given the multipoint SNP genotype data. We then used a database of genotypes of the eight founder strains at 33 million SNP variants (based on a resequencing project at Sanger) to get inferred SNP genotypes for all 500 DO mice at all SNPs. To save computation time, we assumed constant diplotype probabilities within the intervals between SNPs on the GigaMUGA array and took the average of the probabilities for the two endpoints as the probabilities within the interval. We used the founder strains’ SNP genotypes to collapse the 36-state diplotype probabilities to 3-state SNP genotypes. The inferred SNP genotype was that with maximum marginal probability, provided it was > 0.95. Calculations were performed in R (R Core Team 2018) using R/qtl2 (Broman *et al.* 2019).

### Sample mix-ups

For each microbiome sample, we counted the number of reads with each allele at the locations of the 33 million SNPs among the eight founder strains. We omitted SNPs that had more than two alleles among the eight founders. We compared each microbiome sample to each of the 500 genotyped samples and calculated a measure of distance by taking, among reads that overlapped a SNP where the genotyped sample was homozygous, the proportion with an allele that was discordant with the inferred SNP genotype.

### Sample mixtures

A number of the microbiome samples appeared to be mixtures of DNA from two mice. For each microbiome sample, we studied the possibility that it was a mixture between the correctly labeled sample and a single contaminant, and considered each of the other 499 genomic DNA samples as the possible contaminant. For a given microbiome sample and a pair of genomic DNA samples, we counted the number of reads that overlapped a SNP, split according to the SNP allele on the read and the joint SNP genotypes for the pair of samples.

We assumed that the microbiome sample contained a proportion *p* of the contaminant, and assumed a sequencing error rate of *ϵ*, and calculated the expected proportion of reads with an A vs. B allele as a function of the genotypes of the two mice that contributed to the mixture (Supplementary Table S1). With this multinomial model, we then derived maximum likelihood estimates (MLEs) for *p* and *ϵ* by numerical optimization, and we calculated the likelihood ratio test (LRT) statistic for the null hypothesis that there is no contamination (*p* = 0), as twice the natural log of the ratio of the likelihood, plugging in the MLEs of *p* and *ϵ*, to the likelihood evaluated at *p* = 0 with *ϵ* estimated under that constraint.

The LRT values throughout are extremely large, and so to identify potential mixtures, we considered a plot of the LRT vs. the estimated mixture proportion, with a sample considered as a mixture with each of the other possible sample as the contaminant, and looked for a separation between one possible contaminant and the others.

Calculations were performed in R (R Core Team 2018), with the R package R/mbmixture (https://github.com/kbroman/mbmixture).

### Data and software availability

Raw microbiome sequence reads are at the Sequence Read Archive, accession PRJNA744213. The GigaMUGA SNP genotypes are from Keller *et al.* (2018) and are available at https://dodb.jax.org. Founder SNP genotypes are available as a SQLite database at Figshare, doi:10.6084/m9.figshare.5280229.v3.

The processed data and the custom R scripts used for our analyses and to create the figures and tables are at GitHub, https://github.com/kbroman/Paper_MBmixups.

An R package, R/mbmixture, for performing the mixture analysis is available at the Comprehensive R Archive Network (CRAN), https://cran.r-project.org/package=mbmixture, and on Github, https://github.com/kbroman/mbmixture.

## Results

Of the 300 DO mice considered for the microbiome portion of this project, 297 gave usable data, with a median of 39 million reads, counting paired-end reads individually; the vast majority had more than 10 million reads. There was considerable variation in the proportion of reads that were derived from the host. The median was 10.2%, but 25 mice had < 1% and 56 mice had > 25%. There was a strong batch effect, with the second batch of microbiome samples having much lower proportions of reads coming from the host than the other two batches. The reads mapping to the mouse genome spanned all chromosomes, and the absolute counts had a median of 3.9 million, with 78% of samples having > 1 million reads mapping to the mouse genome, and just 6 samples having < 100,000 reads mapping to the mosue genome.

We used high-density SNP genotypes, from the GigaMUGA array, to infer the 36-state diplotypes in each of the DO mice, using a hidden Markov model to calculate the diplotype probabilities conditional on the multipoint SNP data. (See Supplementary Figure S1 for a genome reconstruction of one DO mouse, along with the detailed diplotype probabilities along one chromosome.) We then used a database of the eight founder strains’ genotypes at 32 million autosomal SNPs to infer the corresponding SNP genotypes in the DO mice. The median number of missing genotypes in the imputed data was only 64,466 (0.2%), though there were seven samples with more than a million missing values in the imputed genotypes, including one sample (DO-306) with 14 million missing values in the imputations, though this sample still had 18 million imputed genotypes and was not among the mice with microbiome data.

### Sample mix-ups

In order to identify possible sample mix-ups, we compared each microbiome sample to each genomic DNA sample, by finding sequence reads that overlapped a SNP and counting the number of sequence reads in the six categories defined by the SNP genotype in the genomic DNA sample and the SNP allele on the sequence read. We coded the SNP alleles based on their frequency in the founder strains (A for the major allele and B for the minor allele). At SNPs where a mouse is AA, the microbiome reads should be mostly A, and at SNPs where a mouse is BB, the microbiome reads should be mostly B. We used the proportion of discordant microbiome reads, among those overlapping a SNP where the genomic DNA sample was homozygous, as a measure of distance between the microbiome sample and the genomic DNA sample.

For example, Supplementary Table S2 contains counts of sequence reads from the microbiome sample for mouse DO-360, classified by the SNP genotype of DO-360 and by the allele present in the read. Among the 10.4 million reads that overlapped a SNP where DO-360 had genotype AA, 14.6% had a B allele, and among the 1.2 million reads that overlapped a SNP where DO-360 had genotype BB, 55.6% had an A allele. The distance between microbiome sample DO-360 and genomic DNA sample DO-360 was thus 0.19, the overall proportion of discrepant calls.

If we compare microbiome sample DO-360 to genomic DNA sample DO-370 (Supplementary Table S3), however, we find that only 0.2% of reads overlapping an AA SNP had a B, and only 0.5% of reads overlapping a BB SNP had an A, for an overall distance of 0.002. Similarly, microbiome sample DO-370 is most like genomic DNA sample DO-360. Thus, for samples DO-360 and DO-370, either the genomic DNA samples got swapped or the microbiome samples got swapped. A comparison of RNA-Seq data for these samples (Keller *et al.* 2018) to the SNP genotypes also indicated a sample swap, and so we can infer that the mix-up was in the genomic DNA samples.

We calculated distances between each of the 297 microbiome samples and each of the 500 genomic DNA samples. Figure 1 contains a scatterplot where each point represents one of the microbiome samples, with the x-axis being the distance to the corresponding genomic DNA sample (distance to self), and the y-axis being the minimum distance.

**Figure 1.**
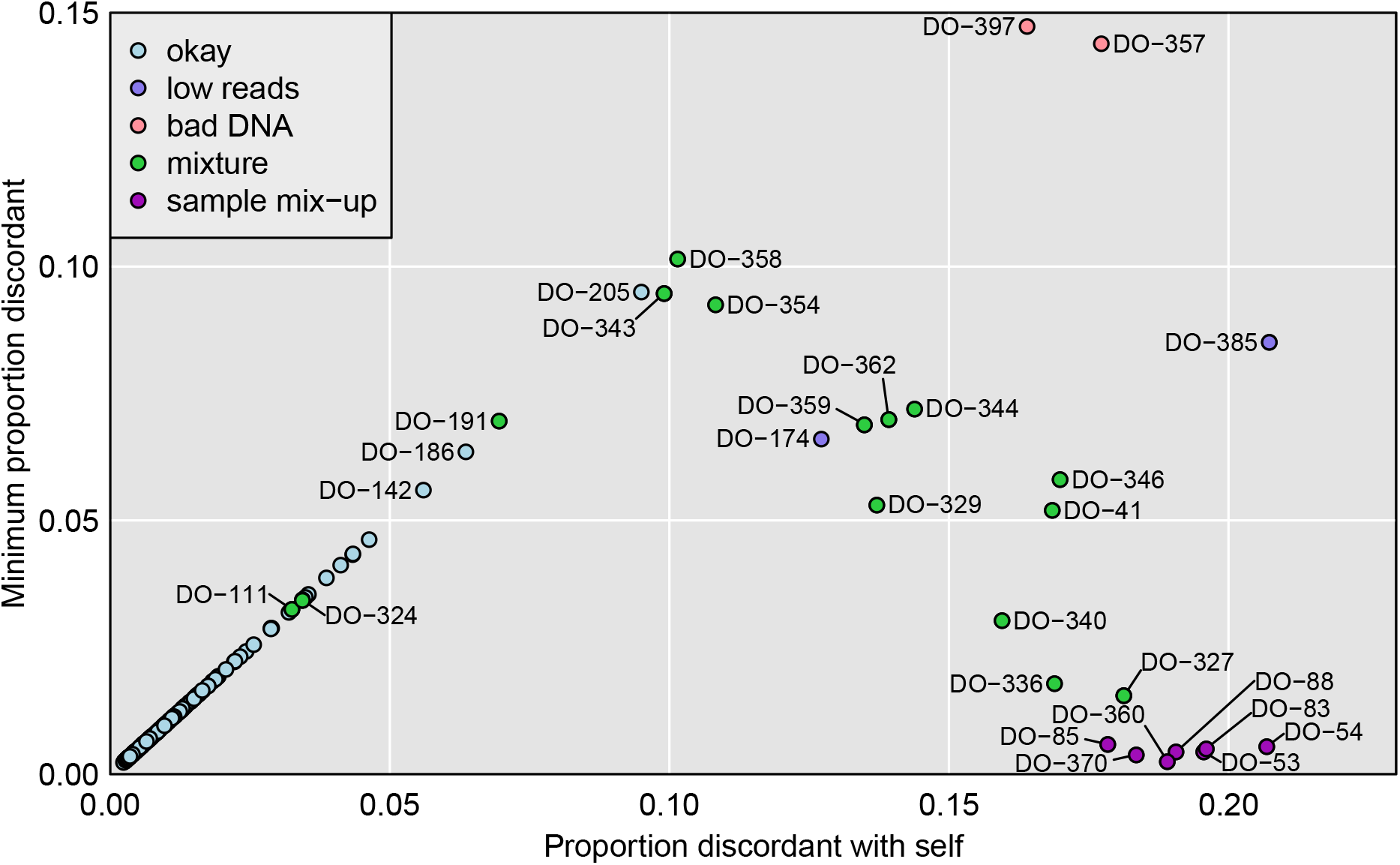
Plot of minimum distance vs. distance to self. Each point is a microbiome sample and distances to genomic DNA samples are measured by taking, among reads that overlapped a SNP where the genotyped sample was homozygous, the proportion with an allele that was discordant with the inferred SNP genotype. The microbiome samples are categorized according to our conclusions about their status.

For microbiome samples along the diagonal, the corresponding genomic DNA sample was the closest, and where this distance is small (lower-left corner), we can conclude that the microbiome sample is correctly labeled. However, as we will see below, a couple of these look to be mixtures, contaminated by a small amount of another sample.

There are seven microbiome samples (colored dark purple in Figure 1) where the distance to the corresponding genomic DNA sample was > 0.15 but the minimum distance to a microbiome sample was < 0.01. These are clear sample mix-ups, and form two pairs and one trio: DO-360/DO-370 (mentioned above), DO-53/DO-54, and DO-83/DO-85/DO-88.

Figure 2 provides further detail on a selected set of microbiome samples, with their distance to each of the 500 genomic DNA samples. As seen in Figures 2A and 2B, samples DO-53 and DO-54 are a clear mix-up (either in the genomic DNA samples or in the microbiome samples) as each microbiome sample is closest to the oppositely labeled genomic DNA sample. DO-101 (Figure 2C) is an example of a correctly labeled sample.

**Figure 2.**
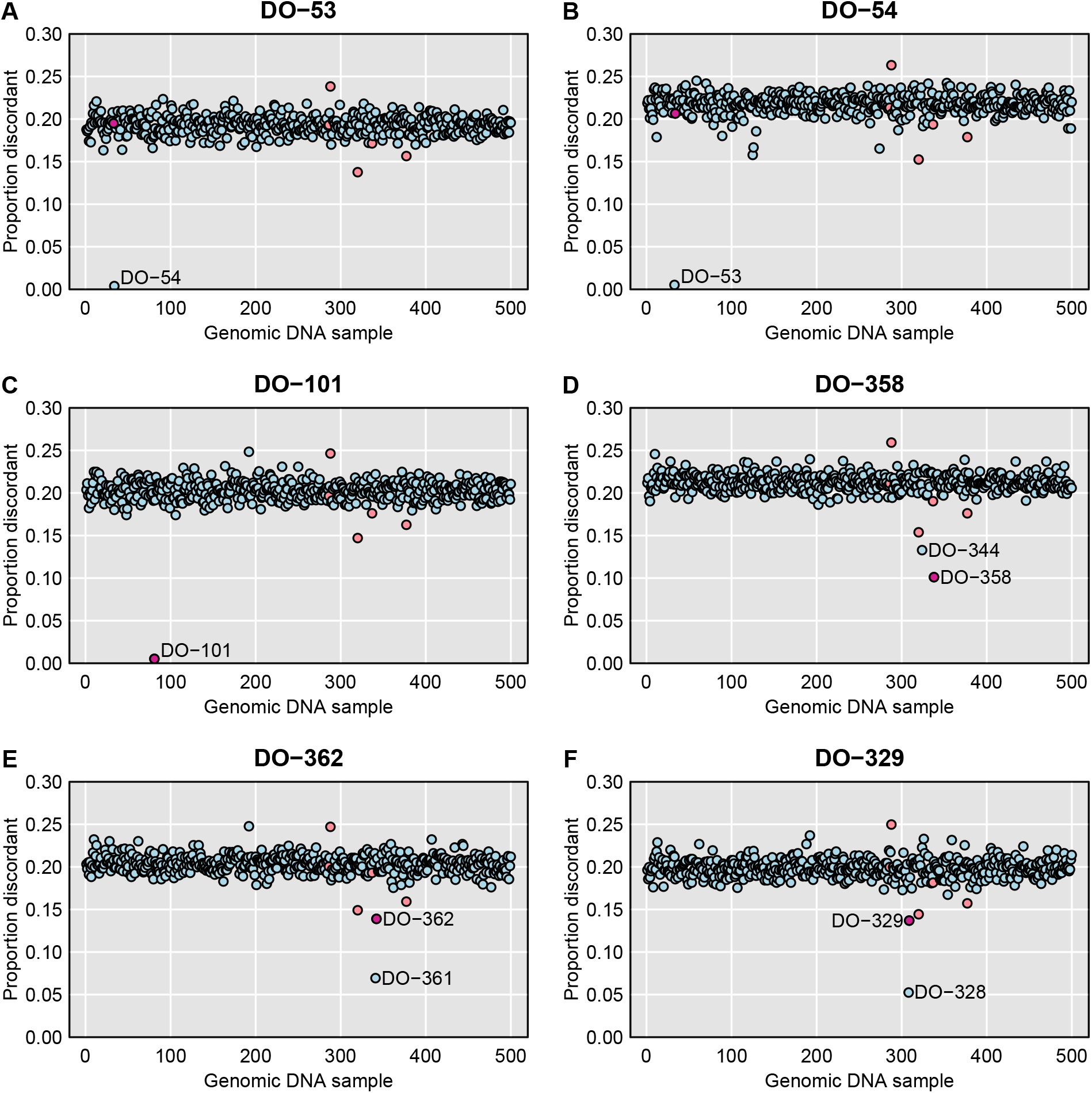
For selected microbiome samples, plots of their distance to each genomic sample, with distance measured by taking, among reads that overlapped a SNP where the genotyped sample was homozygous, the proportion with an allele that was discordant with the inferred SNP genotype. The sample with what should be the correct label is highlighted in dark pink. A set of low-quality DNA samples are highlighted in pale pink.

Additional cases are shown in Supplemental Figure S2. The top row includes the three-way mix-up: microbiome sample DO-83 is closest to genomic DNA sample DO-88, microbiome sample DO-88 is closest to genomic DNA sample DO-85, and microbiome sample DO-85 is closest to genomic DNA sample DO-83.

Returning to Figure 1, there are a number of microbiome samples that are not particularly close to the corresponding genomic DNA sample, nor to any sample. For DO-357 and DO-397 (salmon-colored points), the genomic DNA samples were of low quality. For DO-174 and DO-385 (colored light purple), the microbiome data included only a small number of reads. But another 15 microbiome samples (in green) appear to be mixtures of two samples.

Consider, for example, microbiome sample DO-358. It is closest to the corresponding genomic DNA sample, but at a relatively large distance (0.10), and as seen in Figure 2D, it is relatively close to DO-344 (distance 0.13).

Microbiome sample DO-362 is closer to genomic DNA sample DO-361 than to its own genomic DNA sample, but as seen in Figure 2E, its own sample is second-closest. A similar pattern is seen for microbiome sample DO-329, which is closest to genomic DNA sample DO-328, but its own genomic DNA sample is second-closest. Additional examples are shown in Supplementary Figure S2.

### Sample mixtures

To investigate the possibility that a microbiome sample is a mixture of two samples, we break down the counts of sequence reads by the SNP genotype at two genomic DNA samples. For example, Supplementary Table S4 contains sequence read counts for microbiome sample DO-358, split according to the SNP genotypes at genomic DNA samples DO-358 and DO-344. If the microbiome sample DO-358 contains only DNA from DO-358, then the proportion of reads with the B allele shouldn’t depend on the genotype at DO-344, but in reads overlapping SNPs where both DO-358 and DO-344 are AA, 0.3% have a B, while in reads overlapping SNPs where DO-358 is AA but DO-344 is BB, 40.9% have a B. This is strong evidence for the microbiome sample DO-358 being a mixture of DO-358 and DO-344, and 40.9% is a rough estimate of the proportion of the sample that came from DO-344. (Further, we can estimate the sequencing error rate as 0.3%.)

In Table S5 we consider reads for microbiome sample DO-101, split according to the SNP genotypes at genomic DNA samples DO-101 and DO-102. In contrast to Table S4, we see that the frequency of A reads does not depend on the genotype at DO-102; the DO-101 microbiome sample does not appear to be contaminated by DO-102.

More formally, we fit a multinomial model to the read counts for a microbiome sample, assumed to be a mixture of the corresponding genomic DNA sample plus one other, and derive derive maximum likelihood estimates (MLEs) of the contaminant probability *p* (the proportion of the microbiome sample that came from the second genomic DNA sample) and the sequencing error rate *ϵ*. We further calculate a likelihood ratio test (LRT) statistic for a test of *p* = 0 (that is, the microbiome sample having no contamination).

For microbiome sample DO-358, considered as a mixture of DO-358 and DO-344, we estimate the contaminant probability as 42.0% and the sequencing error rate as 0.3%. The LRT statistic (for testing *p* = 0) is 1.1 ×10^6^.

Figure 3 displays the results for each microbiome sample, considered as a mixture of the correct sample plus each possible other genomic DNA sample, one at a time, as a contaminant. On the y-axis is the LRT statistic for the contaminant sample that gave the largest such statistics, and on the x-axis is the corresponding estimate of the contaminant probability.

**Figure 3.**
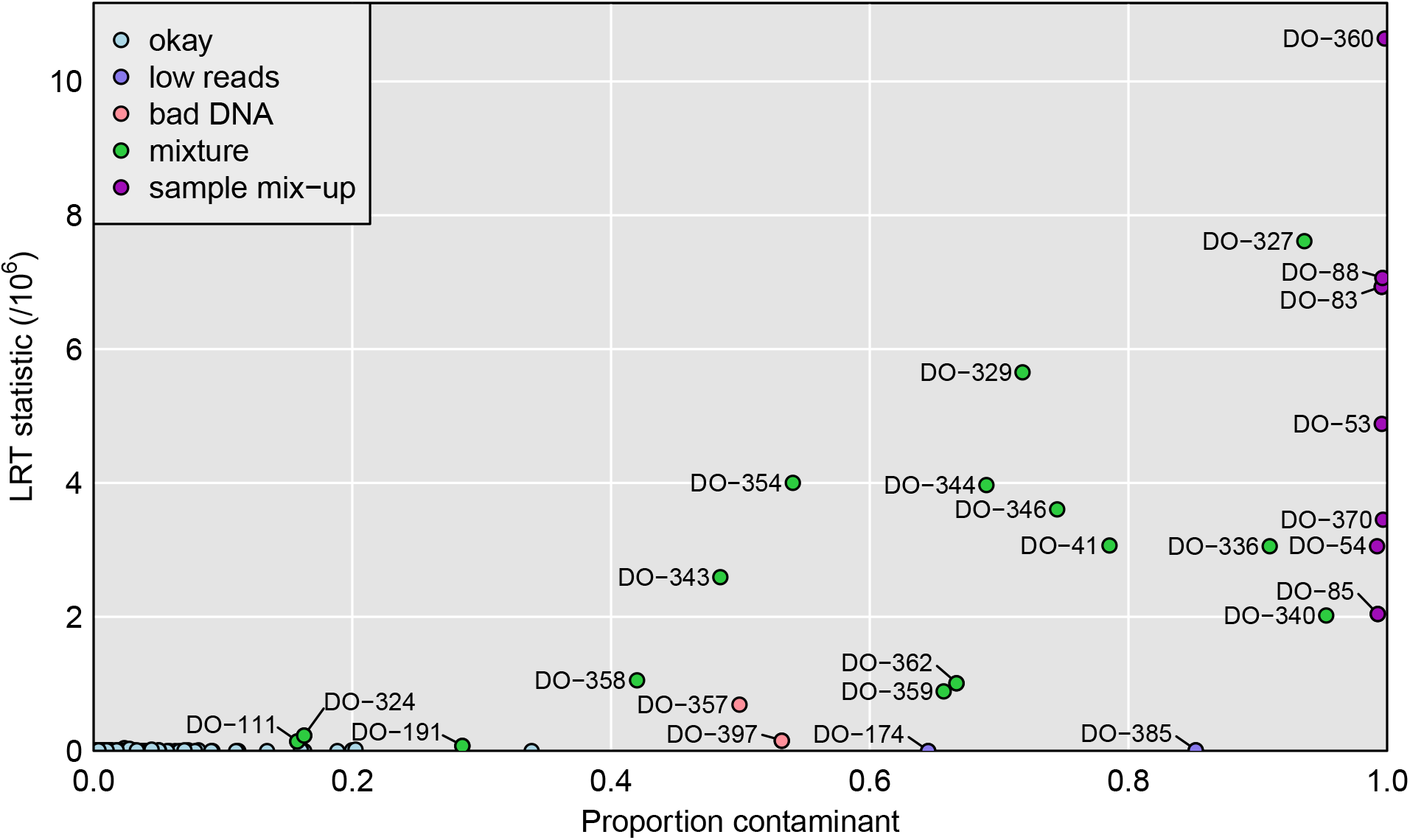
Plot of maximum achieved likelihood ratio test statistic (for the test of no mixture) vs. the estimated proportion from a contaminant, for each microbiome sample. The samples are categorized as in Figure 1, by our conclusions about their status.

The seven sample mix-ups, mentioned previously, are on the far right of Figure 3, with estimated contaminant probability near 1, and with large values for the LRT statistic. There are 15 apparent mixtures, shown in green. Of these, 12 are quite clear, with estimated contaminant probabilities between 42% and 95%, and with LRT statistics > 8 ×10^5^. There are three others (DO-111, DO-191, and DO-324) that are more subtle, with estimated contaminant probabilities of 16–29%, and with LRT statistics of 8–23 ×10^4^.

Figure 4 provides more detailed results for a selected set of samples. Each panel is a scatterplot of the LRT statistic vs. the estimated contaminant probability, for a single microbiome sample considered as a mixture of itself and one other genomic DNA sample, and with the points corresponding to the other 499 genomic DNA samples. In each case, a single possible contaminant stands out. Figures 4A and 4B are for the apparent mix-up of DO-360 and DO-370, and each is estimated to be 100% from the other mouse.

**Figure 4.**
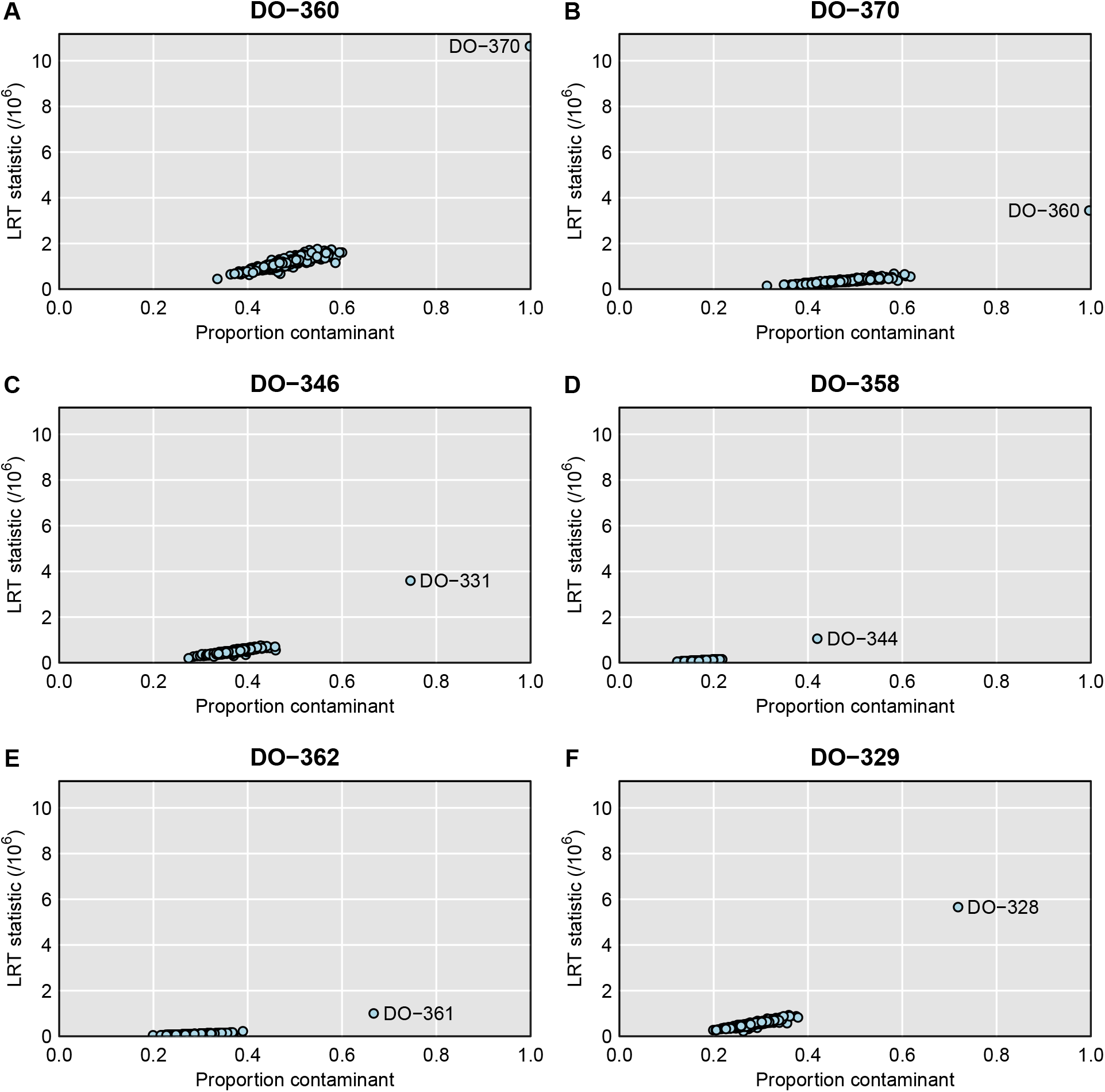
For selected microbiome samples, plots of the likelihood ratio test statistic vs. the estimated proportion contaminated, when considered with each of the genomic DNA samples, one at a time, with the assumption that the microbiome sample is a mixture of the correct sample and that particular contaminant.

Figures 4C–4F are four selected mixtures: DO-346 looks to be 74% from DO-331, DO-358 looks to be 42% from DO-344, DO-362 looks to be 67% from DO-361, and DO-329 looks to be 72% from DO-328. The LRT statistic is a measure of the strength of evidence for the sample being a mixture; it depends both on the contaminant probability as well as total number of sequence reads.

Additional cases are shown in Supplemental Figure S3, including the three more subtle cases (DO-111, DO-191, and DO-324), for which the LRT statistic is a factor of 10 smaller. For these cases, the single-sample analysis to investigate sample mix-ups (see Figure 1) looked okay, but the mixture analysis indicates reasonably strong evidence that these microbiome samples are contaminated. Supplementary Tables S6 and S7 contain the detailed read counts for microbiome sample DO-111 (considered as a mixture with DO-112) and DO-191 (considered as a mixture with DO-146). In each case, the proportion of sequence reads with a B allele clearly depends on the genotype of the second sample.

### More subtle mixtures

In Figure 5, we expand the lower-left corner of Figure 3. This reveals additional microbiome samples where the LRT statistic for contamination is a factor of 10 or 100 smaller, but where there is still compelling evidence for contamination.

**Figure 5.**
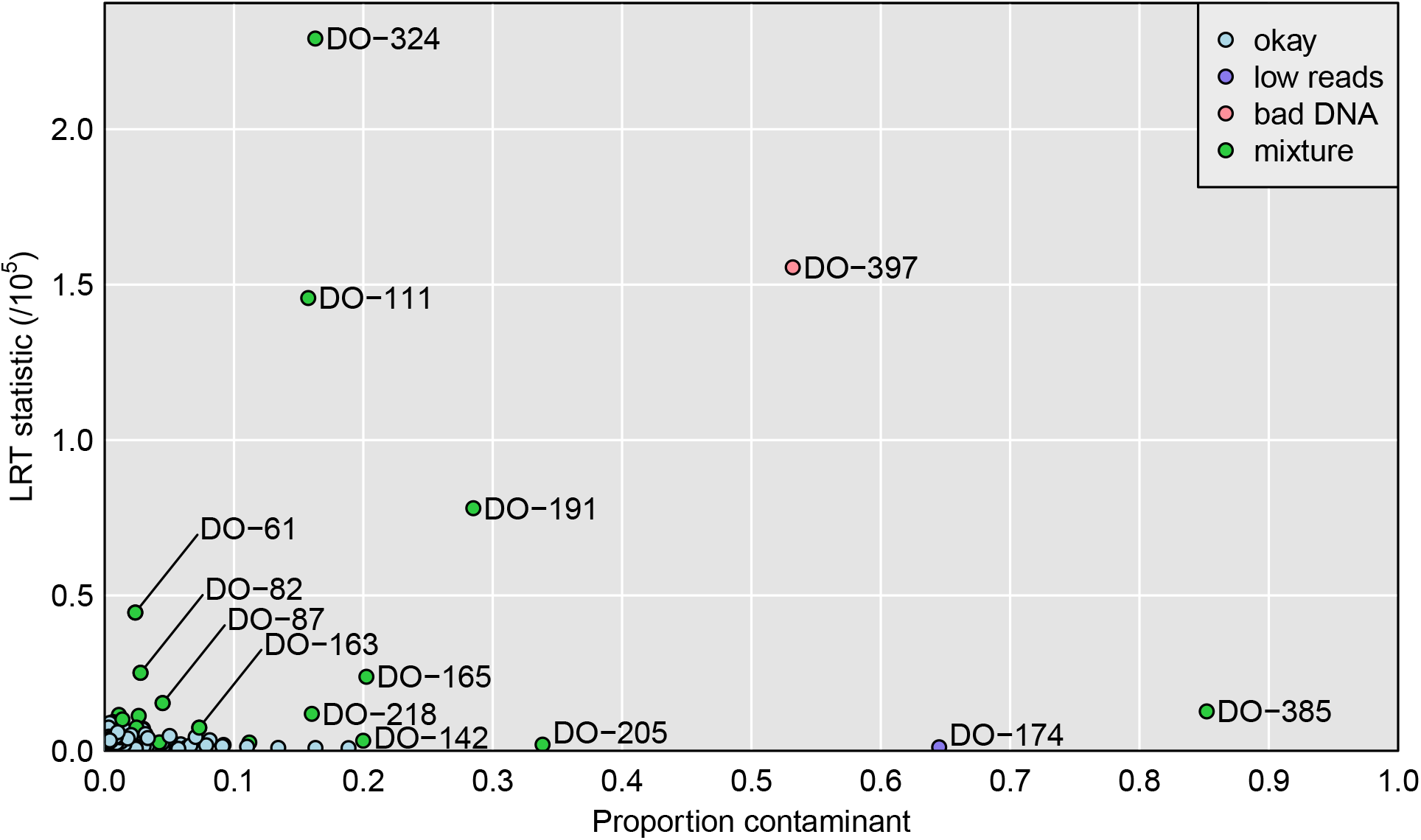
Expansion of the lower-left of Figure 3: Plot of maximum achieved likelihood ratio test statistic (for the test of no mixture) vs. the estimated proportion from a contaminant. The samples are categorized as in Figure 1, by our conclusions about their status.

The minimum LRT statistic across all samples was 279, which would be nominally significant evidence for contamination, but which we believe reflects a weakness of our multinomial model. Before concluding contamination, we consider plots of the LRT statistic vs. the estimated contaminant proportion. Figure S4 displays these plots for 20 selected samples: 17 show clear separation of one potential contaminant from the others, while the other 3 show no such outlier. We consider these 17 samples to be likely contaminated, whereas the other 3 are not.

Among these more subtle sample mixtures is DO-385, which we had previously ruled out as simply having insufficient reads, and DO-205, which showed a high self-distance in Figure 1. Both of these samples have high estimated sequencing error rates (5.7% and 3.6%, respectively).

Many of these subtle mixtures have low estimated rates of contamination, including 10 with estimated contamination proportion < 5%, and with many approaching the estimated sequencing error rate of about 0.6%.

## Discussion

We have described a method for identifying sample mix-ups in metagenomic data for which there are sufficient sequence reads from the host, as well as corresponding SNP genotypes on the hosts. In our study of 500 Diversity Outbred mice, with metagenomic data on 297 of the mice, we identified three sets of sample mix-ups (two pairs and one trio). In a separate aspect of the project (Keller *et al.* 2018), we had generated RNA-Seq data on 300 of the 500 mice, but only 200 were in common with 297 mice for which we have metagenomic data. One of the three sets of sample mix-ups we identified was also seen when the RNA-Seq and SNP data were compared, and so we conclude that the sample mix-up concerned the genomic DNA. For the other two sets of sample mix-ups we identified, the mice were not assayed for mRNA expression, and so we cannot know whether the mix-ups were in the genomic DNA samples or in the microbiome samples.

In the analysis to identify possible mix-ups, we identified a number of microbiome samples that were similar to a pair of genomic DNA samples. We suspected that these were mixtures of a pair of samples (one correct and one contaminant). We derived a method for analyzing such mixtures, which provides an estimate of the proportion of the microbiome sample that comes from the contaminant sample. Among the 297 microbiome samples, 15 had strong evidence of being mixtures of two samples, with estimated contaminant proportions of 16–95% (and with 10 having > 50% from the contaminant sample). An additional 17 samples showed more subtle but still compelling evidence of being mixtures. Many of these had low estimated rates of contamination, including 10 with estimated contamination proportion < 5%.

The basis of our analysis is to identify reads that overlap an informative SNP, and to compare the SNP allele in the read to what would be expected, given the SNP genotype for a particular genomic DNA sample. Our approach thus relies on deep shotgun sequencing of the microbiome samples as well as dense SNP genotypes on the genomic DNA samples. For the latter, we used high-density SNP arrays in the DO mice, in conjunction with a database of SNP genotypes on the eight founder strains, to produce very-high density SNP genotype information in the DO mice. Our approach might be applied to other populations; the two key features that are needed are detailed host genotype data and sufficient metagenomic reads that are derived from the host.

Our approach to identify mix-ups was simplistic: simply counting discrepant reads at homozygous SNPs, pooled across all SNPs. One might instead seek to sufficient metagenomic reads at individual SNPs in order to determine SNP genotypes, but our approach is more expedient. Another alternative would be to perform QTL mapping with microbiome-derived phenotypes and identify sample mix-ups using an approach like that developed for gene expression data (e.g., Westra *et al.* 2011). This would have the advantage of not relying on host-derived metagenomic reads, but would require numerous microbiome QTL with strong effects.

Metagenomic data on microbiome samples provides a more rich understanding of the bacterial functions in the diverse population in the samples, relative to what may be learned from single-locus 16S ribosomal RNA gene sequencing. It also provides the ability to identify and potentially correct sample mix-ups and mixtures in the microbiome samples, which can further strengthen a project’s results.

Samples that appear to be mixtures with high proportions of contamination should likely be excluded from further analyses. But samples with more subtle contamination, say < 5%, may still provide useful data and need not be excluded. The contamination adds to the phenotypic noise for such traits as the abundance of individual bacterial species, or for measures of bacterial diversity. But the contamination, if at a low rate, may be swamped by other sources of variation.

## Acknowledgments

The authors thank the University of Wisconsin Biotechnology Center DNA Sequencing Facility for providing sequencing and support services. They also thank the University of Wisconsin Center for High Throughput Computing in the Department of Computer Sciences for providing computational resources and support. Amelie Baud and an anonymous reviewer provided valuable comments for improvement of the manuscript. This work was supported in part by National Institutes of Health grants R01GM070683 (to K.W.B.), R01DK101573 (to A.D.A.), and R01DK108259 (to F.E.R.), and Postdoctoral Fellowship T15LM007359 (to L.L.T.).

---

**Table S1.** Expected proportion of A microbiome reads, assuming it is a mixture of two genomic DNA samples, as function of the genotypes of the two samples.

| sample 1 genotype | sample 2 genotype | expected proportion of A allele |
| --- | --- | --- |
| AA | AA | $1 - \epsilon$ |
| AA | AB | $(1 - p)(1 - \epsilon) + p/2$ |
| AA | BB | $(1 - p)(1 - \epsilon) + p\epsilon$ |
| AB | AA | $(1 - p)/2 + p(1 - \epsilon)$ |
| AB | AB | $1/2$ |
| AB | BB | $(1 - p)/2 + p\epsilon$ |
| BB | AA | $(1 - p)\epsilon + p(1 - \epsilon)$ |
| BB | AB | $(1 - p)\epsilon + p/2$ |
| BB | BB | $\epsilon$ |

$p$ is the proportion of the microbiome sample coming from genomic DNA sample 2.
$\epsilon$ is the rate of sequencing errors.

**Table S2.** Read counts in microbiome sample DO-360 broken down by observed SNP allele and by the genotype of genomic DNA sample DO-360.

| DO-360 genotype | allele in DO-360 microbiome |  |  |  |
| --- | --- | --- | --- | --- |
|  | A (%) |  | B (%) |  |
| AA | 8,863,572 | (85.4%) | 1,520,169 | (14.6%) |
| AB | 2,870,063 | (72.7%) | 1,075,126 | (27.3%) |
| BB | 671,722 | (55.6%) | 536,010 | (44.4%) |

**Table S3.** Read counts in microbiome sample DO-360 broken down by observed SNP allele and by the genotype of genomic DNA sample DO-370.

| DO-370 genotype | allele in DO-360 microbiome |  |  |  |
| --- | --- | --- | --- | --- |
|  | A (%) |  | B (%) |  |
| AA | 10,324,265 | (99.8%) | 23,256 | (0.2%) |
| AB | 2,083,947 | (51.2%) | 1,986,380 | (48.8%) |
| BB | 5,347 | (0.5%) | 1,117,994 | (99.5%) |

**Table S4.** Read counts in microbiome sample DO-358 broken down by observed SNP allele and by the genotypes of genomic DNA samples DO-358 and DO-344

| DO-358 genotype | DO-344 genotype | allele in DO-358 microbiome |  |  |  |
| --- | --- | --- | --- | --- | --- |
|  |  | A (%) |  | B (%) |  |
| AA | AA | 2,394,215 | (99.7%) | 6,050 | (0.3%) |
| AA | AB | 869,613 | (79.5%) | 224,483 | (20.5%) |
| AA | BB | 103,036 | (59.1%) | 71,332 | (40.9%) |
| AB | AA | 686,970 | (71.8%) | 269,447 | (28.2%) |
| AB | AB | 297,500 | (51.4%) | 280,958 | (48.6%) |
| AB | BB | 55,982 | (29.9%) | 131,111 | (70.1%) |
| BB | AA | 73,727 | (42.9%) | 98,257 | (57.1%) |
| BB | AB | 47,000 | (21.9%) | 167,513 | (78.1%) |
| BB | BB | 542 | (0.5%) | 117,802 | (99.5%) |

**Table S5.** Read counts in microbiome sample DO-101 broken down by observed SNP allele and by the genotypes of genomic DNA samples DO-101 and DO-102

| DO-101 genotype | DO-102 genotype | allele in DO-101 microbiome |  |  |  |
| --- | --- | --- | --- | --- | --- |
|  |  | A (%) |  | B (%) |  |
| AA | AA | 3,664,076 | (99.6%) | 14,305 | (0.4%) |
| AA | AB | 1,161,383 | (99.5%) | 6,187 | (0.5%) |
| AA | BB | 153,501 | (99.3%) | 1,067 | (0.7%) |
| AB | AA | 651,287 | (52.0%) | 600,434 | (48.0%) |
| AB | AB | 378,800 | (51.8%) | 352,828 | (48.2%) |
| AB | BB | 155,967 | (51.2%) | 148,703 | (48.8%) |
| BB | AA | 3,088 | (1.6%) | 185,825 | (98.4%) |
| BB | AB | 3,210 | (1.3%) | 240,712 | (98.7%) |
| BB | BB | 2,162 | (1.0%) | 217,882 | (99.0%) |

**Table S6.** Read counts in microbiome sample DO-111 broken down by observed SNP allele and by the genotypes of genomic DNA samples DO-111 and DO-112

| DO-111 genotype | DO-112 genotype | allele in DO-111 microbiome |  |  |  |
| --- | --- | --- | --- | --- | --- |
|  |  | A (%) |  | B (%) |  |
| AA | AA | 1,378,006 | (99.6%) | 5,121 | (0.4%) |
| AA | AB | 352,914 | (92.7%) | 27,682 | (7.3%) |
| AA | BB | 25,268 | (85.8%) | 4,180 | (14.2%) |
| AB | AA | 337,647 | (59.3%) | 231,369 | (40.7%) |
| AB | AB | 202,368 | (52.2%) | 185,606 | (47.8%) |
| AB | BB | 33,524 | (44.3%) | 42,090 | (55.7%) |
| BB | AA | 21,171 | (17.0%) | 103,638 | (83.0%) |
| BB | AB | 9,266 | (8.8%) | 96,292 | (91.2%) |
| BB | BB | 768 | (1.0%) | 77,959 | (99.0%) |

**Table S7.** Read counts in microbiome sample DO-191 broken down by observed SNP allele and by the genotypes of genomic DNA samples DO-191 and DO-146

| DO-191 genotype | DO-146 genotype | allele in DO-191 microbiome |  |  |  |
| --- | --- | --- | --- | --- | --- |
|  |  | A (%) |  | B (%) |  |
| AA | AA | 333,088 | (99.4%) | 2,144 | (0.6%) |
| AA | AB | 132,182 | (85.7%) | 22,011 | (14.3%) |
| AA | BB | 19,120 | (73.8%) | 6,772 | (26.2%) |
| AB | AA | 77,327 | (67.0%) | 38,093 | (33.0%) |
| AB | AB | 43,599 | (54.6%) | 36,208 | (45.4%) |
| AB | BB | 9,291 | (37.7%) | 15,329 | (62.3%) |
| BB | AA | 5,343 | (28.3%) | 13,556 | (71.7%) |
| BB | AB | 3,368 | (14.7%) | 19,534 | (85.3%) |
| BB | BB | 88 | (0.6%) | 13,517 | (99.4%) |

**Figure S1.**
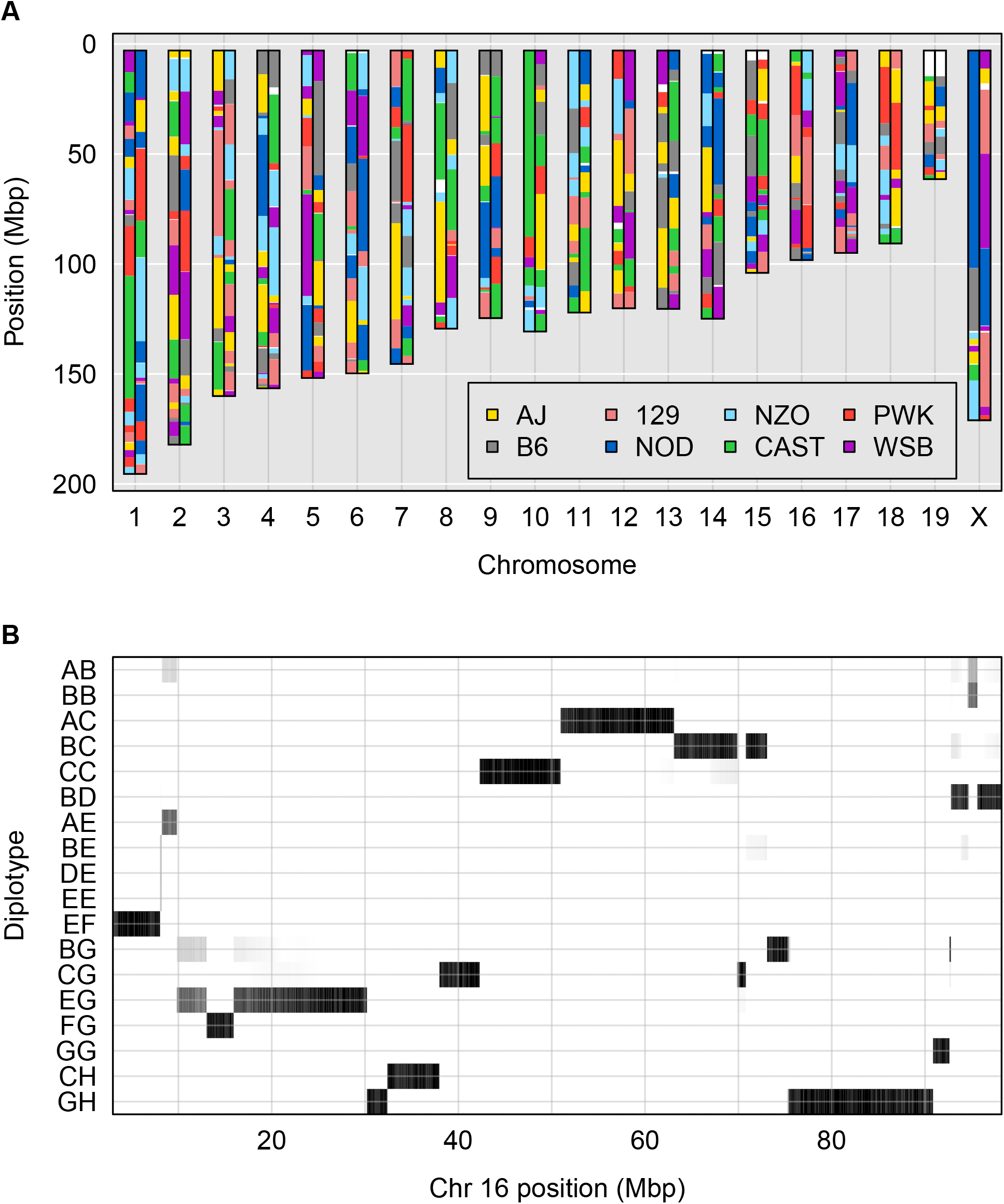
Illustration of genome reconstruction. **A**: Inferred diplotype across the genome for mouse DO-359, with an arbitrary choice of phase. (For example, if there is a region of homozygosity, the haplotypes above can be swapped relative to the haplotypes below.) White indicates unknown (no diplotype had probability >0.5). **B**: Heat map of the diplotype probabilities for mouse DO-359 along chromosome 16. Only diplotypes that achieved probability > 0.25 are shown.

**Figure S2.**
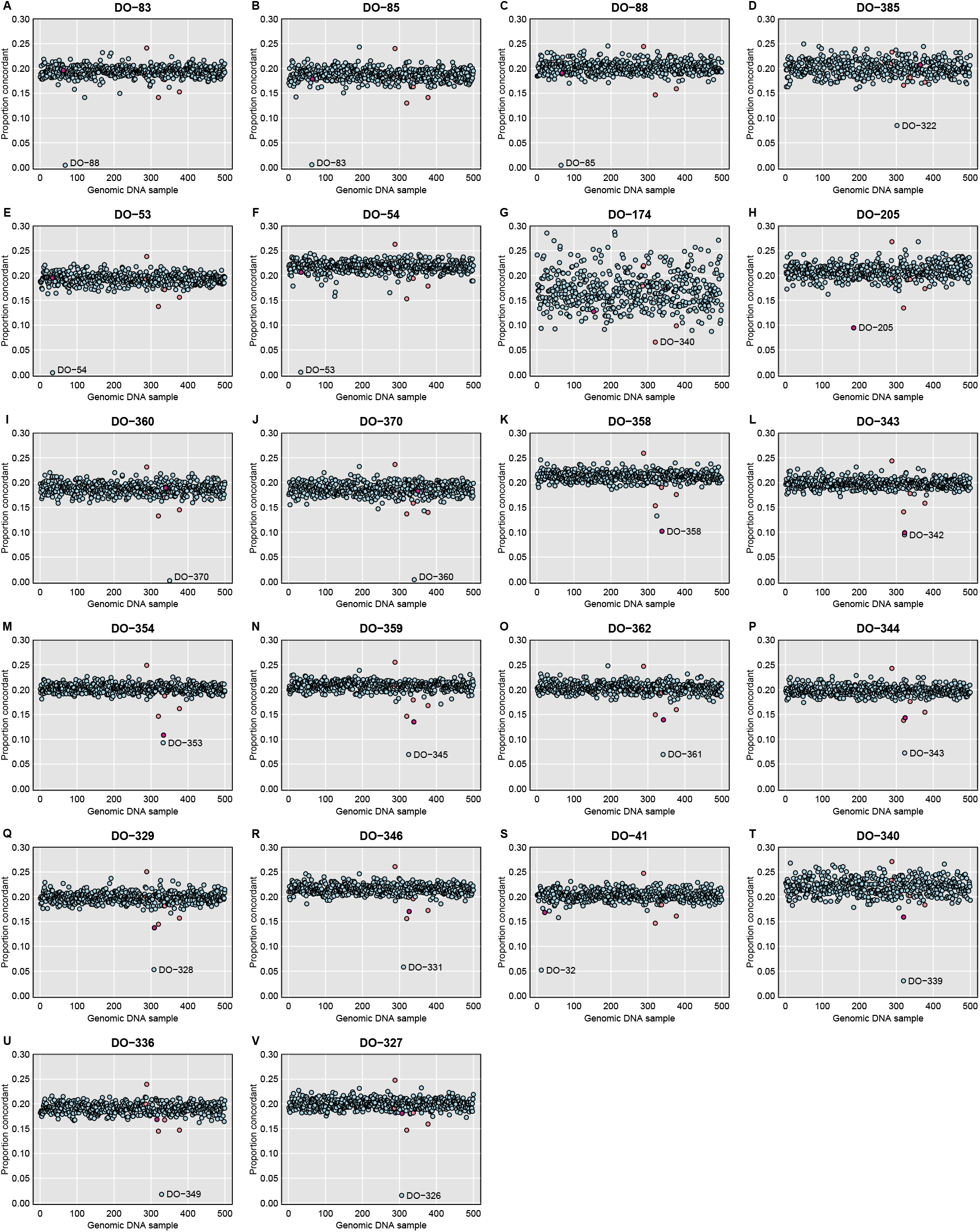
For selected microbiome samples, plots of their distance to each genomic sample, with distance measured by taking, among reads that overlapped a SNP where the genotyped sample was homozygous, the proportion with an allele that was discordant with the inferred SNP genotype. The sample with what should be the correct label is highlighted in dark pink. A set of low-quality DNA samples are highlighted in pale pink.

**Figure S3.**
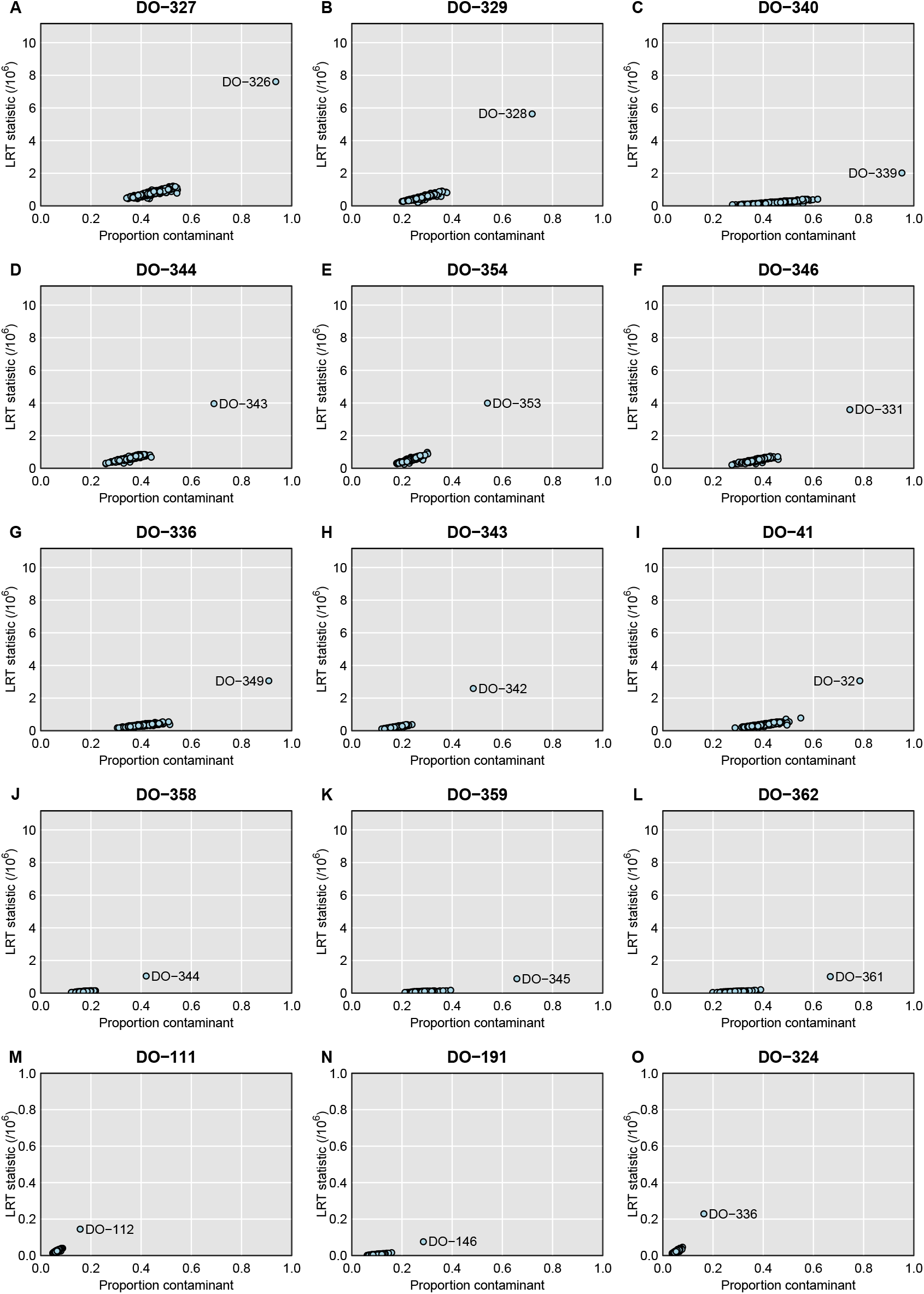
For selected microbiome samples, plots of the likelihood ratio test statistic vs. the estimated proportion contaminated, when considered with each of the genomic DNA samples, one at a time, with the assumption that the microbiome sample is a mixture of the correct sample and that particular contaminant. Note the change in the y-axis limits in the bottom row of panels.

**Figure S4.**
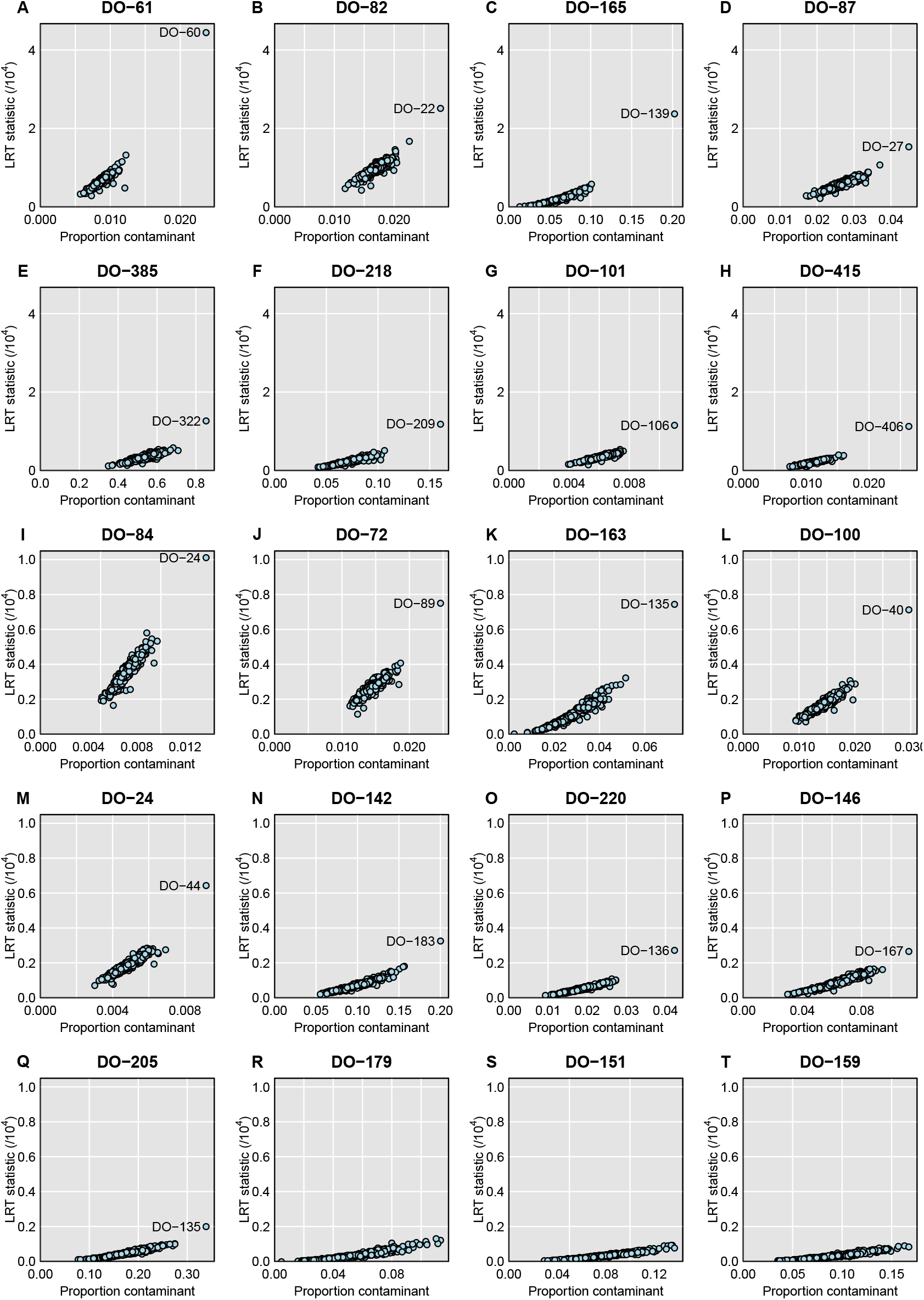
For further selected microbiome samples, plots of the likelihood ratio test statistic vs. the estimated proportion contaminated, when considered with each of the genomic DNA samples, one at a time, with the assumption that the microbiome sample is a mixture of the correct sample and that particular contaminant. Note the change in the y-axis limits in the bottom three rows of panels.

